# Systematic and comprehensive benchmarking of an exome sequencing based germline copy-number analysis pipeline to detect clinically relevant CNVs

**DOI:** 10.1101/579755

**Authors:** Ramakrishnan Rajagopalan, Jill Murrell, Minjie Luo, Laura K. Conlin

## Abstract

**Purpose:** Detecting germline copy-number variants (CNVs) from exome sequencing (ES) is not a standard practice in clinical settings owing to several reasons concerning performance. We comprehensively characterized an ES-based CNV pipeline and developed frameworks for minimizing false-positives and assess the reproducibility.

**Methods:** We used a cohort of 387 individuals with both clinical chromosomal microarray (CMA) and ES data available to estimate the initial performance by comparing CNVs from both platforms. A modification of the default workflow was performed to reduce the number of false positives and the reproducibility of the CNVs was assessed using an iterative variant calling process.

**Results:** The default pipeline was 93% sensitive with a high false-discovery rate of 44%. The modified workflow had a higher sensitivity of 96% while reducing the total number of CNVs identified and improving the false-discovery rate to 11.4%. With the modified workflow, we demonstrated a 100% validation rate for the CNVs identified in the *STRC*, a challenging gene to ascertain by short-read NGS. The exome-based pipeline was 100% sensitive for clinically-relevant, rare variants (including single exon deletions), and was reproducible.

**Conclusion:** We demonstrate with our modified workflow and the benchmarking data that an exome-based CNV detection pipeline can be reliably used to detect clinically-relevant CNVs.

## Introduction

Exome Sequencing (ES) is the current standard of care diagnostic tool for identifying a molecular cause(s) in individuals with suspected Mendelian disorders^1^. Identifying single nucleotide variants (SNVs) and small insertion/deletions (indels) from next-generation sequencing (NGS) data have been well studied and characterized^2,3^. Success has been elusive to date to detect copy-number variants (CNVs) from NGS data with the same confidence as the SNVs/indels. Chromosomal microarrays (CMA) arrays are still the preferred methodology for detecting genome-wide CNVs^4^. CNV detection using ES is not currently a routine clinical test, likely due to the overwhelming inconsistencies among different methods^5–7^ and the lack of a high-quality reference for CNVs from ES data. Most of the algorithms for CNV detection from ES data use the depth of coverage of exome targets under the assumption that the read depth is linearly correlated with the underlying true copy number at any given locus. However, the read-depth in ES is known to be extremely variable and influenced by several factors such as sample batching, GC-content, PCR duplication bias, targeted depth, sequencing efficiency and mappability^8,9^. These factors make it difficult to differentiate between technical artifacts and the real signal for a true copy-number change. Also, detecting CNVs in polymorphic regions of the genome is challenging as the methods for ES-based CNV are estimating copy-number relative to the average copy-number of the control samples.

While the sensitivity of several software tools for ES-based CNV has been published, reports of false-discovery rate and the reproducibility^7,10^ are limited. Quality and performance standards for a clinical pipeline is set at the highest level possible as it has direct implications on patients’ health and disease management. In spite of the availability of several computational tools to detect CNVs from ES^11–14^, clinical labs have been slow to adopt the incorporation of CNV detection in their ES. However, some of the recent reports support the argument for copy-number detection from ES in a clinical setting^15–17^.

In this work, we used a cohort of samples with clinical CMA and ES data to create a dataset of high-quality true-positive CNVs from CMA and comprehensively characterize the CNVs identified from ES data. The ES-based CNV workflows used in this paper are provided in the *Figure 1*. We addressed some of the rarely discussed issues such as false-discovery rate, false-negatives, and reproducibility. In addition, we propose a modified analysis workflow to reduce false-positives originating from the repetitive regions of the genome. Finally, we show that ES data can be used reliably for detecting clinically-relevant CNVs with high sensitivity in a reproducible manner for use in a clinical diagnostic setting.

**Figure 1.** The original (A) and the modified (B) exome-based CNV detection and validation workflow. 1) Filter for CNVs that overlap at least 10 probes in the array and one bait in the exome design, 2) Remove false-positives in the CMA data by manual review after initial comparison to refine the baseline true-positive CNVs 3) Remove highly polymorphic regions before calculating the false-discovery rate 4) Exons with low mean mappability score were excluded prior to calling CNVs

## Materials and Methods

### Cohort

A cohort of 395 individuals who were referred to the Genomic Diagnostic Laboratory (GDL) at the Children’s Hospital of Philadelphia (CHOP), Philadelphia PA for genetic testing and had both clinical CMA and ES data was collected for this study. We used a larger cohort including an additional 1,585 individuals with both affected individuals referred for genetic testing and their relatives as controls for read-depth normalization and targeted *STRC* testing (n=1,972). All of the ES data were produced using the Agilent SureSelect V5 plus target capture kit (Agilent Technologies, Santa Clara, CA) and the aligned BAM files were produced using the workflow described elsewhere^18^. All of the ES data were generated at the same sequencing center in 112 different batches over 2 years. These samples were analyzed as part of a process improvement program within the GDL at CHOP.

### CNV from Exome sequencing (ES)

We used a custom CNV detection pipeline and analysis workflow using the R package ExomeDepth^19^ with default parameters and exon definitions provided along with the package. ExomeDepth creates a custom reference sample set for every sample by choosing the most correlated control samples from a larger cohort of controls and identifies CNVs based on relative read-depth of the test sample compared against this custom reference panel. Further details of this method are described elsewhere^19^. Mean number of CNVs per sample identified in the entire cohort were calculated. On average each sample had 164 CNVs with the default pipeline and the standard deviation was 27. Samples with number of CNVs identified greater than two standard deviations from the mean were excluded as outliers. There were 8 such outlier samples that were excluded and the final sample size was 387 (207 males and 187 females). Summary statistics of the CNVs identified from the ES are provided in the Table 1(A) All coordinates are in GRCh37.

**Table 1.** Description of the workflow presented in this paper

|  |  |  |
| --- | --- | --- |
| **Step 1** | **Defining the initial gold standard** | Filter for CNVs that overlap at least 10 probes in the array and one bait in the exome design |
| **Step 2** | **Estimate the initial sensitivity** | Estimate initial sensitivity by comparing CNV from ES and CMA and review false-negatives in the exome |
| **Step 3** | **Refine the gold-standard CNVs and sensitivity** | Remove false-positives in the CMA data and recalculate sensitivity |
| **Step 4** | **Estimate the false discovery rate** | Manually verify CNVs called by exome that overlap at least 10 probes and not identified by the array |
| **Step 5** | **Analyze the false positives** | Analyze false-positives to identify factors that influence CNV calling from ES data |
| **Step 6** | **Modify the workflow** | Use mappability filter to exclude regions of high homology and reduce false-positives |
| **Step 7** | **Estimate the reproducibility** | Estimate reproducibility of ES-based CNV workflow by performing tens of thousands of experiments with randomized control cohorts |

### CNVs from SNP arrays

SNP array data were generated using three different platforms, Illumina CytoSNP 850k chip v1.0 (n=142) and v1.1 (n=165), Illumina Omni1 Quad (n=28) and Illumina Quad610 (n=52). Poorly performing probes and probes within highly polymorphic CNV regions (>10% internal cohort) were removed prior to CNV calling. CNVs from SNP arrays were called using CNV-workshop^20^ or PennCNV^21^. CNVs involving fewer than 10 probes were excluded, as the rate of false positives increases with fewer number of probes^22^. We excluded any CNVs called in the chromosome Y due to lack of coverage on both the array and exome capture. CNVs identified from the Quad610 arrays were in hg18 genome build and the liftOver^23^ tool from UCSC was used to convert the genomic coordinates to GRCh37. CNVs that were split in the genome build were excluded prior to any analysis.

### Defining gold standard high-quality true-positive CNVs from clinical SNP arrays

We considered several factors while defining the gold standard baseline true positive dataset for CNVs from CMA, as it dictates the resulting sensitivity of the ES. In order to perform a reasonable comparison between these platforms, we first limited the CNVs from arrays to have at least one coding exon and at least one bait in the ES design, and overlap at least 10 SNP probes in order to minimize the number of false positive and false negatives from the CMA data set. In cases where we detected a CNV in the array but not in the ES, we manually reviewed the raw data from CMA to determine if the call was a false negative in the ES or a false positive in the CMA. This process was mainly performed to refine the baseline true positive calls as even the list of CNVs overlapping at least 10 probes is likely to have its own false positive and false negative rates. Failing to remove the putative false positives from the CMA will deflate the actual sensitivity of the ES.

In our validation cohort with the SNP array data, there were a total of 9,033 CNVs prior to exclusion. Applying the exclusion criteria resulted in 634 CNVs (589 in the autosomes and 45 in the chromosome X). These 634 CNVs formed our initial true positive dataset for comparison against the CNV calls from ES data. Fifty-five of these CNVs were not detected in the ES data. These discrepant calls were manually reviewed, and 41 were confirmed to be true positives in the array, seven were false positives in the array, and seven were ambiguous. The fourteen CNVs that were either not present or ambiguous in the CMA were excluded and the revised baseline true positive dataset contained 620 CNVs (576 autosomal and 44 chromosome X variants). Details of this filtering cascade are provided in Table 2 and the summary statistics of the baseline true-positive CNVs are provided in Table 3. The final list of all the true positive CNVs is provided in *Supp. Table 1*.

**Table 2.** Characteristics of the CNVs identified from the CMA and the ES data

| Characteristics of the baseline truth CNVs defined by the CMA |  |  |
| --- | --- | --- |
|  | Deletion | Duplication |
| Number of CNVs | 234 | 386 |
| Size | 2.1kb to 3.1 Mb (mean = 162kb, median = 71kb) | 3.3kb to 5.4 Mb (mean = 243kb, median = 107kb) |
| # of SNP probes | 10 to 2,258 (mean = 61, median = 20) | 10 to 774 (mean = 62, median = 29) |
| # of exons overlapping the CNV | 1 to 462 (mean = 13, median = 6) | 1 to 341 (mean = 20, median = 10) |
| Small CNVs (< 4 exons) | 52 (22%) | 94 (24%) |
| Reported CNVs | 34 (15%) | 24 (6%) |
| Characteristics of the CNVs identified from ES data |  |  |
|  | Deletions | Duplications |
| Number of CNVs | 30,727 | 24,948 |
| Size | 2 bp to 2.5 Mb (mean = 13kb, median = 974bp) | 2 bp to 9.5 Mb (mean = 16kb, 1.4kb) |
| # of exons | 1 to 470 (mean = 4, median = 2) | 1 to 981 (mean = 4, median = 2) |
| # of CNVs per individual | 91 | 73 |

**Table 3.**
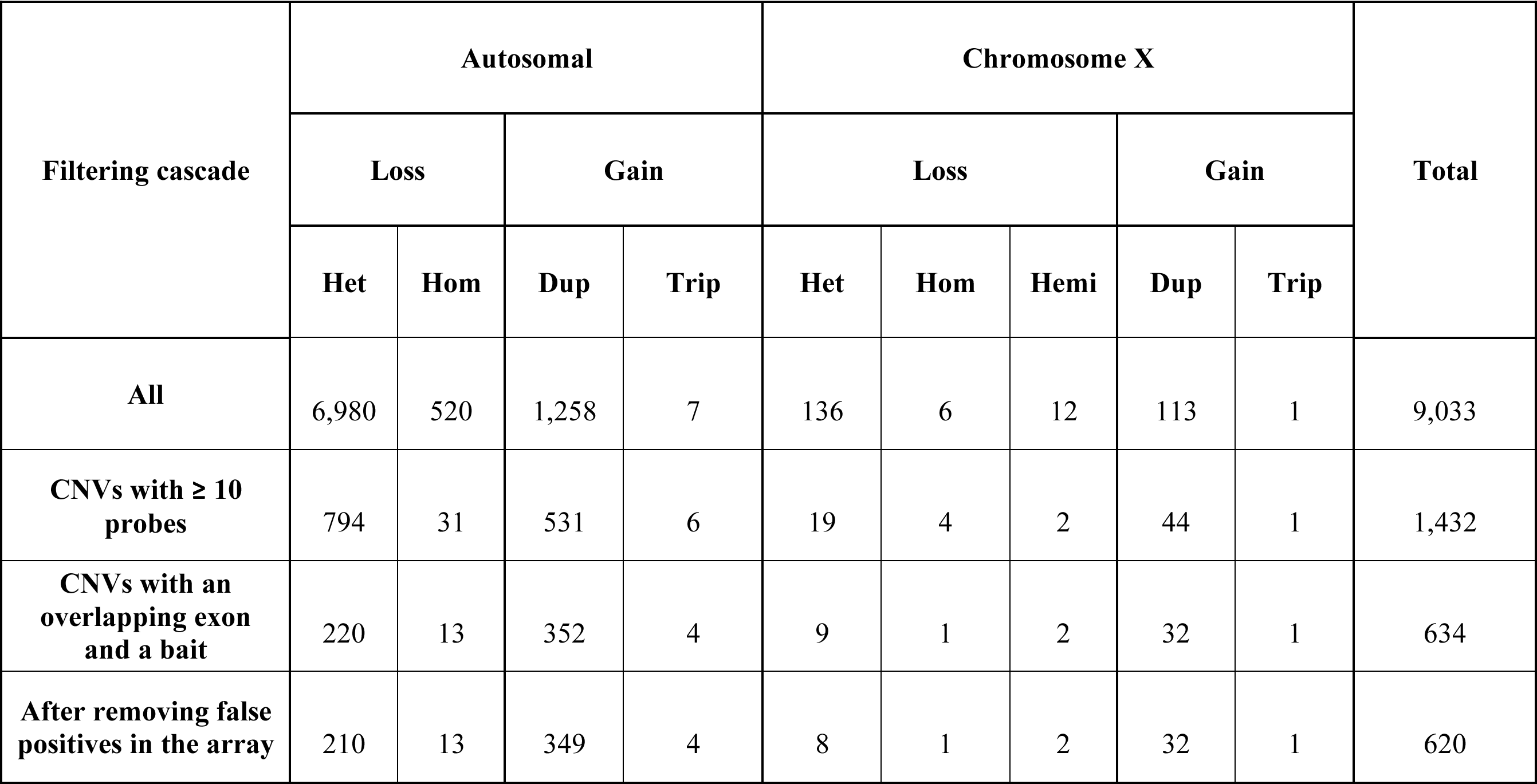
Details of the filtering cascade for the chromosomal microarray data to create a list of baseline true positive CNVs

### Estimating false discovery rate of CNV from ES

To estimate the false discovery rate of the ExomeDepth pipeline, we reviewed CNVs called from ES data from the most recent 124 samples with a clinical genome-wide array analysis of the same chip type (Illumina CytoSNP 850k v1.1). From these samples, there were 385 CNVs from the ES data that overlapped at least 10 probes in the CMA design. Forty-two percent of these calls originated from two known extremely polymorphic regions, the immunoglobulin-like receptor region in chr19 (chr19:55,236,714-55,367,367) and the HLA region in chr6 (chr6:32,549,335-32,709,302). We did not review the CNVs from these two regions as they are known to be highly polymorphic and challenging in both platforms. The remaining 225 CNVs were compared the CMA data and a subset of the variants not identified by the default CMA pipeline (n = 122) were manually reviewed by looking at the log R ratio, B-allele frequency and genotype clustering of all the SNP array probes overlapping the CNV. To understand the factors that influence the false-positives, we annotated each CNV with exon-level allele frequencies from DGV, number of overlapping CNV calls from the entire cohort with the same and opposite copy-number state, mean alignability, mean mappability and percent overlap with repeat regions in the genome^24^.

### Modifying the existing workflow

We used the 35-mer mappability score^25^ from the UCSC genome browser (link) and excluded any exon with a mean mappability score less than or equal to 0.75 prior to the variant calling. We excluded 8,527 out of the total 190,340 unique exons we used in the initial experiment prior to calling the CNVs. We applied the modified regions of interest to both the 387 samples, plus a larger cohort of 1,585 samples to assess the effect of this filter on the clinically-relevant gene *STRC*, which is known to have a pseudogene with 99.6% sequence identity.

After the modification of the workflow, we used the same cohort described in the previous section (n=124) and repeated the protocol to estimate the estimate false-discovery rate. We manually reviewed the remaining 149 CNVs to estimate the false-discovery rate after the modification of the workflow.

### Reproducibility of the ExomeDepth pipeline

To test the effect of the control cohort on reproducibility, we ran 1,000 iterations of our pipeline for each of the 387 samples, along with a subset of 200 controls randomly chosen from a larger control cohort of 1,585 samples. We kept track of 1) the number of reference samples selected, 2) batch from which the control sample was sequenced, 3) correlation of the coverage between the test and the reference samples, and 4) the CNV calls produced. We also specifically looked at 59 clinically reported variants in 53 individuals for their reproducibility over the 1,000 iterations.

### Validation of CNVs using Droplet-Digital PCR (ddPCR)

All diagnostic CNVs and a subset of *STRC* CNVs were chosen for confirmation using PCR across the breakpoints, droplet digital PCR (ddPCR), or long-range PCR, using standard protocols used in the clinical laboratory^26^.

## Results

Using a cohort of 387 samples, we validated an exome-based CNV detection pipeline using the R package ExomeDepth^19^ for use in a clinical setting, by comparing the results against data from a high-quality set of true positive CNVs from CMA. We chose ExomeDepth after a pilot study comparing other software such as XHMM^11^, CoNIFER^12^, and CODEX^13^.

ExomeDepth was the most sensitive of the algorithms tested and the union of results from all three were no better than ExomeDepth alone (*data not shown*). We estimated the true positive and false discovery rates and implemented a modified bioinformatics pipeline to reduce the total number of CNVs identified and the number of false positives. Finally, we estimated the reproducibility of 59 clinically reported CNVs in 53 patients by running 1,000 experiments per sample along with randomly chosen subsets of controls from a larger cohort.

### True positive rate

Overall, the default ES pipeline had a 94% true positive rate for deletions (219/233) and 93% for duplications (361/ 387). In the exomes, 96% percent of the true positives were identified as single contiguous CNVs and the rest as two CNV calls. The ES pipeline identified all homozygous deletions (n=14), hemizygous deletions (n=2) and triplications (n= 5). Sensitivity for the autosomes was 94% (539/576) compared to the chromosome X at 91% (40/44). Clinically reported variants in the CMA data were detected with 100% sensitivity (52/52). Summary of sensitivity rates from the default workflow for various CNV classes are provided in the Table 4(A).

**Table 4.** Sensitivity of the ExomeDepth pipeline

| **True-positive rate** | **Deletions** | **Duplications** |
| --- | --- | --- |
| Overall | 94% (219/234) | 93% (360/386) |
| Heterozygous deletions | 93% (203/ 218) |  |
| Homozygous deletions | 100% (14/14) |  |
| Hemizygous deletions | 100% (2/2) |  |
| Duplications |  | 93% (355/381) |
| Triplications |  | 100% (5/5) |
| Autosomal | 94% (210/223) | 94% (329/353) |
| Chromosome X | 82% (9/11) | 94% (31/33) |
| Reported CNVs | 100% (34/34) | 100% (25/25) |
| CNVs overlapping 1 exon | 82% (23/28) | 69% (24/35) |
| CNVs overlapping 2 exons | 69% (11/16) | 89% (32/36) |
| CNVs overlapping 3 exon | 80% (4/5) | 100% (18/18) |
| CNVs overlapping >3 exons | 98% (180/184) | 96% (287/298) |

**Table 5.** New diagnoses made by the ES pipeline that were previously not

| **ID** | **Genomic coordinates (hg19)** | **Gene** | **Size (bp)** | **CNV** | **# of exons** | **# of SNPs in the CMA** | **Disease inheritance** |
| --- | --- | --- | --- | --- | --- | --- | --- |
| Patient 1 | chr4:123,976,639-123,989,201 | *SPATA5* | 12,562 | Heterozygous deletion | 2 | 3 | Recessive |
| Patient 2 | chr12:116,457,804-116,461,341 | *MED13L* | 3,537 | Heterozygous deletion | 1 | 3 | Dominant |
| Patient 3 | chr2:149,210,869-149,219,983 | *MBD5* | 9,114 | Heterozygous deletion | 1 | 3 | Dominant |
| Patient 4 | chr6:33,405,980-33,409,266 | *SYNGAP1* | 3,286 | Heterozygous deletion | 5 | 4 | Dominant |
| Patient 5 | chr3:191,888,248-192,126,012 | *FGF12* | 237,765 | Duplication | 4 | 123 | Dominant |

There were 41 CNVs from the CMA that were not identified by our exome pipeline. Of these 41 false negatives (37 autosomal and 4 chromosome X), 15 were heterozygous deletions and 26 were duplications. Size of the false negatives ranged from 2kb to 260kb for the deletions (1 – 20 exons) and 16kb to 1.2Mb (1 – 29 exons) for the duplications. Thirty-three of the 41 false negatives were in highly polymorphic or segmental duplicated regions in clinically irrelevant areas of the genome. Thirteen false negative CNVs involved first or last exons. A single-exon, intragenic duplication in the *PRKN* gene was not identified by the ES pipeline. Details of the CNVs missed by the exome pipeline are provided in the *Supp. Table 2*.

### False discovery rate

In order to determine the false discovery rate from ES, we analyzed CNVs identified by the ES pipeline that overlapped at least 10 probes in the CMA that should have been theoretically identified by the CMA. Of the 225 variants reviewed, 103 were identified by the default CMA pipeline and remainder of the 122 CNVs manually reviewed, 22 were determined to be real in the CMA and one was determined to be real but with an opposite copy-number state in a polymorphic region, likely due difference in the average copy-number state in the controls.

Together we found forty-four percent (44%) of the CNVs were deemed to be false-positives in the ES data. Details of the CNVs identified from the exomes and not by the CMA are provided in the *Supp. Table 3*.

### Modified ES workflow

Manual review of the false-positive CNV calls from ES data suggested that a large number of those calls were identified in regions with high homology elsewhere in the genome or regions with low sequence complexity. To mitigate this effect, we used a mappability filter to exclude exons that have difficulty in mapping short-read NGS data^27^ prior to CNV calling. To assess the effect of the mappability filter, we compared the average number of variants identified per sample with the modified workflow against the results from the initial analysis. With the mappability filter in place, the average number of calls identified reduced from 164 per sample (114 to 278 with a median of 160) to 50 per sample (32 to 166 with a median of 49). Comparing to the true positive calls, we observed a higher sensitivity of 96% compared to the default workflow. We repeated the protocol to estimate the false-discovery rate by manually reviewing the 149 CNVs and found that 17 of them were false-positives in the ES data leaving the false-discovery rate at 11.4%.

To quantify the effect of the mappability filter on CNVs identified from highly homologous regions of the genome, we ran this modified workflow on a larger cohort of 1,972 samples. Further, we chose to focus on the *STRC* gene, as it is one of the most common causes of autosomal recessive nonsyndromic hearing loss and our clinical lab has a validated protocol to confirm the copy-number variants in this gene. The *STRC* gene has a pseudogene that is 99.6% identical to the protein-coding gene, with the first 15 of the 29 exons identical between the two genes. We excluded the exons of *STRC* with a mappability score less than 0.75, leaving 3 exons in the modified workflow. With the original workflow, there were 953 CNVs in 731 individuals (37% of the samples) and 147 individuals had both a deletion and duplication identified in *STRC*. With the modified workflow, we identified a total of 76 *STRC* CNVs (29 deletions and 47 duplications) in 76 individuals (4% of the samples). Of these, we performed confirmatory testing on 6 duplications and 24 deletions using ddPCR, 2 single-exon deletions by long range PCR, and 3 multi-exon deletions by CMA. We were able to confirm all multi-exonic deletions and duplications, and the single-exon deletions were found to be associated with gene conversion events involving exon 26 and exon 24. Sensitivity of the modified ExomeDepth workflow is provided in Table 4(B). Details of the validation data are provided in the *Supp. Table 4*.

### Reproducibility

Previous reports suggest that, among other factors, the choice of the control cohort affects the CNVs identified from ES data. To understand the effect of the choice of the control cohort, we estimated reproducibility of our pipeline by running 1,000 iterations for every sample with 200 random controls and counted the number of times the clinically reported CNVs were called. Over these 387,000 experiments, the average number of calls per sample across iterations was 60 for the autosomes (median = 50) and 2 for the X chromosome (median = 1.4). The number of batches from which these reference samples came ranged from 3 to 14 (mean = 9). There was a total of 3,204 CNVs in the cohort that were identified in all 1,000 iterations, with a mean of 8 CNVs per individual. Of these CNVs which met criteria to be called by the array (n=221), 179 were also identified by the CMA (81%). To resolve the discrepant calls (n=42), we manually reviewed a subset of CNVS (n=14) and confirmed that 12 out of the 14 were actually false negatives in the CMA. Of all the CNVs we were able to confirm with array data, 190 out of 192 CNVs were real and reproducible. All the clinically-reported variants including 52 CNVs from the CMA and 5 CNVs from the ES, were identified in all 1,000 experiments for those samples.

### New diagnoses

In the process of the validation, several previously unrecognized clinically significant CNVs were discovered. All of the probands included in the cohort (n = 351) were referred to exome sequencing after a negative or inconclusive SNP array. We made a total of 5 new molecular diagnoses in 5 individuals who had negative or inconclusive results with the SNV/indel only pipeline or previously not identified in the SNP array (overlapping fewer than 10 probes in the SNP array) or not being in a known disease-causing gene at the time of reporting.

Where possible, we were able to determine the exact breakpoints for three deletions from the chimeric reads in the exome data and validated these using Sanger sequencing across the breakpoints. The details of the new diagnoses are provided in the *Supp. Table 5*. These pathogenic CNVs included 1 to 9 exons, in both autosomal dominant and recessive disease associated genes.

## Discussion

Detecting CNVs from ES data is perceived as challenging for clinical use as many previously published reports suggested high false positive rates and low sensitivity. In this work, we used a cohort of 387 individuals to systematically and comprehensively benchmark the ability to detect CNVs from exome sequencing data using the ExomeDepth pipeline. Prior benchmarking efforts have attempted to use data from multiple orthogonal platforms and algorithms for estimating sensitivity and specificity from ES data^7,27,28^. Performance metrics such as sensitivity and false-discovery rate rely solely on the baseline true positive set. Factors influencing such comparisons should be considered carefully before defining the true positive set as the resulting performance metrics may be misleading. Our diagnostic lab uses a 10-probe threshold for CMA data as CNVs overlapping fewer probes are known to have a higher rate of false positives^22^. ES has poor sequencing efficiency in GC-rich regions, in segmental duplication regions, and in regions with low sequence complexity^8,9^. Also, the sequencing is based on a target design which may or may not include regions of the genome that is captured by an alternative technology. Taking the inherent limitations of both the ES and CMA technology into account allowed for the maximization of the clinical utility of the ES data. Several algorithms exist for detecting CNVs from ES and there is a trade-off between the true positive rate (detecting the true CNVs) and false discovery rate (detecting false positives). For use in a clinical setting, one requires the highest sensitivity and lowest false discovery rate possible for a given platform.

We defined a high-quality baseline true positive set for CNVs using data from CMA and found that ES is 93% sensitive for both deletions and duplications using the default settings of the ExomeDepth pipeline. The analysis of CNVs missed by ES (false negatives) revealed that 80% of them were in highly polymorphic or segmental duplicated regions and 31% of them involved first or last exons. First exons are known to be GC rich and GC-content greatly influences the depth of coverage^11^. However, there is no prior evidence associating last coding exons with non-uniform depth of coverage during exome sequencing. Analysis of false positives showed a similar trend with 89% located in overlapping regions that are polymorphic in the general population, of low sequence complexity, or of segmental duplications. The ExomeDepth pipeline coupled with a mappability threshold for including exons before calling the CNVs reduced the number of calls to one third which reduced the burden of downstream analysis and validation. In addition, we were able to detect CNVs in the clinically-relevant and difficult regions, such as *STRC*, with a 100% validation rate.

Using this pipeline, we were able to make new diagnoses in five individuals who had previous negative CMA tests, demonstrating the utility of this assay for small (1 exon to multi-exon CNVs), intragenic CNVs below the clinical reporting threshold for CMA. The results from the ExomeDepth pipeline were 100% reproducible for clinically reported variants with control datasets generated from over 112 batches over a period of 2 years, even though sample batching is known to be strongly correlated with the variability in the depth of coverage observed in ES data^11^.

Despite our positive results, some challenges still exist. For example, it is challenging to determine the exact copy-number state in the highly variable regions of the genome and these regions pose a challenge irrespective of the technology used. A large number of true positives and larger cohorts with homogeneous samples may be required to optimize CNV calling in such polymorphic regions. Detecting CNVs in genes with near identical homologs elsewhere in the genome is an intractable problem when using short-read sequencing data. These regions often result in both false positive and false negative CNV calls. Using an exon-level mean mappability score threshold helps in reducing the false positives but it also excludes some clinically relevant genes completely (e.g. *SMN1* and *SMN2*). On the other hand, this mappability filter still allows the inclusion of exons with partial regions of poor mappability, resulting in numerous false positives. The 100% validation rate we observed for the *STRC* gene deletions and duplications after using the mappability-based filter is encouraging and further work is warranted to understand the effect on similar regions elsewhere in the genome. In spite of being able to detect the clinically reported CNVs 100% of the time in our reproducibility experiments, we found that the choice of controls and the batch in which the controls were sequenced had an effect on the total number of CNVs identified. Only a small subset of variants per sample were identified across all 1,000 iterations and manual verification of these CNVs suggest that they have a higher likelihood of being real. While the number of times a variant is reproduced with different control cohorts may indicate the robustness of the call, it is also likely that technical artifacts are also reproducible. It is important to note that all of our controls are generated in the same sequencing facility with the same protocol. Using data produced from different sequencing protocols, target captures, and sequencing centers could affect the reproducibility and produce erroneous results. Further experimentation is warranted to utilize this information along with other quality metrics for better ranking of likely true positive variants. Validation of clinically-relevant CNVs using an orthogonal method is important before reporting to the patients as ES-based CNV detection is still relative to the control cohort used. However, maintaining good practices in creating a control cohort, a validation pipeline with a variety of known variants types and size, and stringent quality control before clinical correlation will reduce the burden of validations using orthogonal methods.

In summary, our work demonstrates the ability to detect CNVs from ES data in a reliable and reproducible manner in a clinical setting. Integrating CNVs in a clinical workflow may help in finding molecular diagnoses for unresolved patients with one pathogenic variant (SNV/indel) in an autosomal recessive disease gene, and increase the overall diagnostic rate. We expect targeted NGS to be used in diagnostics for a considerable amount of time given the lower cost, focused approach and reduced burden on downstream analysis compared to genome sequencing. Thus, it is important to continue to invest in resources and refine the existing tools for making ES a better diagnostic test overall.

## Table legends

Table 1. Characteristics of the CNVs identified from the ES data with the default workflow (A) and the modified workflow (B)

Table 2. Details of the filtering cascade for the chromosomal microarray data to create a list of baseline true-positive CNVs

Table 3. Characteristics of the CNVs identified from the CMA

Table 4. Sensitivity of the ExomeDepth pipeline with the default workflow (A) and the modified workflow (B)

## Acknowledgements

This work is supported by the National Institutes of Health grant R01-HG009708 and U01-HG006546. We would like to acknowledge the Division of Genomic Diagnostics at the Children’s Hospital of Philadelphia for providing the samples and data for this study. We acknowledge the members of the Bioinformatics group at the Division of Genomic Diagnostics for making the computational resources available. We acknowledge Jorune Balciuniene PhD, Zhiqian Fan MS, and Heather Pearce, BS of the Genomic Diagnostic Laboratory at the Children’s Hospital of Philadelphia for their help in validating the STRC variants.

## Author contributions

RR, LKC designed the study, performed the experiments, analyzed the data and wrote the manuscript. JM analyzed the data, validated the CNVs, discussed the results and edited the manuscript. ML designed validation studies, discussed results and critically reviewed the manuscript.

---

Supp. Table 1: List of true positive CNVs and the exome validation status

Supp. Table 2: List of CNVs missed by the exome (false negatives)

Supp. Table 3: List of CNVs that overlapped 10 SNPs in the CMA and reviewed for determining false-discovery rate

Supp. Table 4: Validated STRC calls

Supp. Table 5: New diagnoses made by the ES pipeline that were previously not reported

